# Augmenting Signaling Pathway Reconstructions

**DOI:** 10.1101/2020.06.16.155853

**Authors:** Tobias Rubel, Anna Ritz

**Affiliations:** Department of Biology, Reed College, Portland, OR, US; Department of Philosophy, Reed College, Portland, OR, US

**Keywords:** Systems biology, signaling pathways, graph algorithms, pathway reconstruction

## Abstract

Signaling pathways drive cellular response, and understanding such pathways is fundamental to molecular systems biology. A mounting volume of experimental protein interaction data has motivated the development of algorithms to computationally reconstruct signaling pathways. However, existing methods suffer from low recall in recovering protein interactions in ground truth pathways, limiting our confidence in any new predictions for experimental validation. We present the Pathway Reconstruction AUGmenter (PRAUG), a higher-order function for producing high-quality pathway reconstruction algorithms. PRAUG modifies any existing pathway reconstruction method, resulting in augmented algorithms that outperform their un-augmented counterparts for six different algorithms across twenty-nine diverse signaling pathways. The algorithms produced by PRAUG collectively reveal potential new proteins and interactions involved in the Wnt and Notch signaling pathways. PRAUG offers a valuable framework for signaling pathway prediction and discovery.

## 1 Introduction

Signaling pathways describe the series of molecular interactions that occur in response to a certain stimulus, which mediates the expression of relevant genes. Understanding the specific reactions that occur within pathways is a fundamental question in molecular systems biology, and through decades of research we now have some understanding of how cells grow, proliferate, and die. Many of these reactions are stored in databases that characterize these pathways such as NetPath [1], KEGG [2], Reactome [3], and dozens of others. Biologists have extensively used these databases to help determine which pathways are activated or inhibited in high-throughput gene expression experiments. Despite the promise of pathway database resources, we know that they are incomplete. The same pathway can be represented very differently across databases, and they often have little overlap among the proteins and interactions. There are also new discoveries about protein membership and interactions involved in canonical signaling pathways that are not in these databases yet. Further, pathways in non-model systems or under-studied pathways after often missing from these resources.

In the past decade, there have been two emerging solutions for improving pathway databases. First, pathways have been integrated into resources that attempt to capture the current knowledge base of signaling. Repositories such as WikiPathways [4] and PathwayCommons [5] now contain thousands of pathways comprised of millions of interactions by aggregating information from other databases. WikiPathways also offers a community-driven platform, allowing researchers to submit and curate pathways [4].

While these resources are useful for exploration, they are not intended to make predictions about *new* proteins and interactions that may be associated with a pathway of interest. Another area of signaling pathway research develops new methods that can predict new players in signaling pathways of interest. Many of these approaches generate new predictions by integrating protein-protein interaction data with gene [6–11] or protein [12–15] expression. Other approaches work to remove biologically implausible predictions [16, 17]. Here, we will focus on methods that use protein-protein interaction data to predict new proteins and molecular interactions involved in canonical signaling pathways.

### 1.1 Network-based Pathway Reconstruction

Protein-protein interactions can be modeled as graphs where the nodes stand in for proteins and the edges represent a (possibly directed) interaction between two proteins. These graphs, called *interactomes*, can be built from experimental dataset repositories and can be weighted by confidence in the interaction, functional relationship, or tissue [16, 18–24]. Interactomes provide a background set of plausible interactions for consideration in a pathway of interest, and network-based methods generally have been successful with amplifying the signal of interacting proteins [25].

We are interested in predicting new proteins and interactions involved with a particular pathway of interest. The **Pathway Reconstruction Problem** is summarized as the following [19]: Given an interactome, a set of receptors in a specific pathway, and a set of transcriptional regulators in the pathway, recover the intermediate proteins and interactions responsible for transmitting the signal from the receptors to the transcriptional regulators. The pathway-specific receptors and transcriptional regulators may come from proteins in existing pathway databases [19, 26] or from experimental data [7–9, 11].

### 1.2 Contributions

We present the Pathway Reconstruction AUGmenter (PRAUG), a higher-order function which maps any pathway reconstruction method to an augmented method that improves pathway reconstruction performance. PRAUG is designed based on the observation that pathway reconstruction methods typically perform well when predicting the proteins in a pathway. PRAUG takes as input a pathway reconstruction method and provides a method which uses a traversal on the input method’s predicted nodes to explore protein interactions. Despite PRAUG’s simplicity, this augmentation improves the protein interaction accuracy of almost any pathway reconstruction method that is used to seed the algorithm. Thus, PRAUG can serve as a framework to boost the performance of any existing method for the Pathway Reconstruction Problem. We highlight PRAUG’s generalizability by running it on an interactome of 612516 weighted interactions to recover 29 diverse human signaling pathways from the NetPath pathway database [1], using 6 state-of-the-art pathway reconstruction methods as seeds. We highlight the improved pathway reconstructions using a case study of the Wnt and Notch pathways.

## 2 Methods

We first describe PRAUG and the pathway reconstruction methods used as input to PRAUG. We then describe the interactome and pathway datasets, and details about assessment.

### 2.1 PRAUG

We are given a potentially directed interactome *G* = (*V, E*), which may have weights *w*_*uv*_ for every edge (*u, v*) ∈ *E*. A pathway *P* = (*V*_*P*_, *E*_*P*_) can be described as a subgraph of *G* (e.g. *V*_*p*_ ⊆ *V* and *E*_*P*_ ⊆ *E*). Pathway *P* includes a set *S* ⊆ *V*_*P*_ of receptors and a set *T* ⊂ *V*_*P*_ of transcriptional regulators (TRs). A solution to the Pathway Reconstruction Problem connects the *sources S* to the *targets T* through the interactome *G*.

Many existing pathway reconstruction methods offer a solution to the Pathway Reconstruction Problem. Here, we focus on the space of methods 𝕄 that take an interactome *G*, a set of sources *S* and targets *T*, and potentially other user-defined parameters which we denote {*}. These methods return a subgraph of *G*. Examples of such methods are outlined in Section 2.2. An instance of a method ℳ ∈ 𝕄 will return a subgraph *H* of *G*:

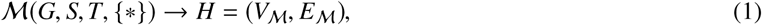

where *V*_ℳ_ ⊆*V* and *E*_ℳ_ ∈*E*. The edges of *H* are often ranked by the order in which they were found by ℳ, but may also be unranked.

PRAUG transforms a method ℳ ∈ 𝕄 into another method 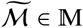. An instance of 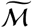 will return a subgraph 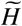 of *G*:

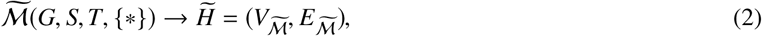

where 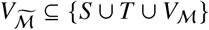 and 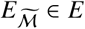. The PRAUG-augmented counterpart of ℳ works as follows:

1. Run ℳ (*G, S, T*, {*}) and return a subgraph *H* = (*V*_ℳ_, *E*_ℳ_).
2. Define a node set *X* = {*S* ∪ *T* ∪ *V*_ℳ_}.
3. Introduce a super source node *σ* to *G* and add edges from *σ* to every node v ∈ *S*.
4. Perform an unweighted depth-first traversal on *G* starting from *σ*, taking only nodes in *X*. Terminate when no new edge can be traversed from *σ* to nodes in *X*.
5. Return the traversed edges as a subgraph 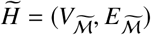, with edges sorted by their traversal order.

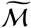 uses the nodes predicted by ℳ as a scaffold to find edges that connect predicted nodes. Some of these edges may not have been discovered by the original ℳ, which may potentially include known interactions for a particular pathway. Since the sources *S* and targets *T* depend on the specific pathway *P* that we wish to reconstruct, we parameterize the methods by the pathway *P* instead of *S* and *T*. We also drop the interactome *G* from the parameterization for simplicity, leaving the methods parameterized as ℳ(*P*, {*}) and 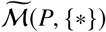.

### 2.2 Pathway Reconstruction Methods

We use six different pathway reconstruction methods as inputs for PRAUG (Table 1).

**Table 1.** Pathway Reconstruction Methods.

| Method Name | Abbrev. | Parameters |
| --- | --- | --- |
| PathLinker [19] | PL | number of shortest paths $k$ |
| ResponseNet [9, 10] | RN | sparsity parameter $\gamma$ |
| BowTieBuilder [26] | BTB |  |
| Prize Collecting<br>Steiner Forest [7, 8] | PCSF | terminal prize $p$<br>edge reliability $b$<br>dummy edge weight $\omega$<br>degree penalty $g$ |
| Random Walk with<br>Restarts (insp. by [11]) | RWR | teleportation probability $\alpha$<br>flux threshold $\tau$ |
| Shortest Paths | SP |  |

#### PathLinker (PL) [19]

This method computes the *k* shortest paths from any source *S* to any target *T* in a weighted, directed interactome. PathLinker uses an *A*^*^ speedup to the classic Yen’s *k*-shortest loopless paths algorithm [19]. PL requires a number of shortest paths *k* ∈ ℤ^+^ (by default *k* = 500), and edges are ranked in increasing order by the first path in which they appear.

#### ResponseNet (RN) [9, 10]

This method formulates the pathway reconstruction problem as a network flow algorithm, and presents a linear program to find a subgraph that balances outgoing flow from the sources and incoming flow to the targets. RN requires a sparsity parameter *γ* ∈ ℝ^+^ that penalizes flow through multiple sources (by default *γ* = 20). The unranked set of edges with positive flow are considered the predicted pathway reconstruction.

#### BowTieBuilder (BTB) [26]

This method iteratively connects sources to targets using short paths that aim to determine an hourglass, or “bowtie” structure in the pathway reconstruction. The method terminates when as many sources and targets as possible have been added to the graph. The unranked set of edges are considered the predicted pathway reconstructionxs.

#### Prize Collecting Steiner Forest (PCSF) [7, 8]

This method places node prizes on the sources and targets in a weighted (but undirected) interactome, and uses a message-passing algorithm to identify a forest (set of trees) that connect prizes while simultaneously minimizing the edge costs [8]. We convert the directed, weighted interactome *G* into an undirected graph by converting any bidirected edge between nodes *u* and v to an undirected edge with the minimum cost of (*u*, v) and (v, *u*); the remaining directed edges simply become undirected. In addition to a prize *p* ∈ ℝ^+^, the method also has a parameter *b* ∈ ℝ+ that controls the tradeoff between including more prizes and taking higher-cost edges, a weight ω ∈ ℝ^+^ for dummy edges, and a degree penalty g ∈ ℝ^+^. Default parameters are *b* = 1,ω = 5, and g = 3. The unranked set of edges in the forest are considered the predicted pathway reconstruction.

#### Random Walk with Restarts (RWR)

Inspired by TieDIE [11], we implemented a random walk with restarts using the following procedure. First, we run a random walk with restarts from the sources *S*, teleporting to sources uniformly at random. We then reverse the edges of the graph and run a random walk with restarts from the targets *T*, teleporting to targets uniformly at random. For a node *v* ∈ *V*, let the forward visitation probability be denoted as *p*_*F*_ (*v*) and let the backward visitation probability be denoted as *p*_*B*_(*v*). We calculate the combined flux *f* (*u, v*) for an edge (*u, v*) as

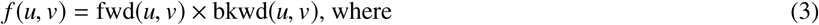

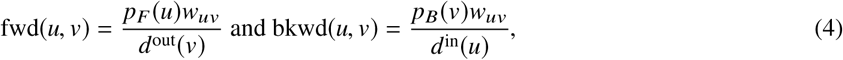

where *d*^in(*v*)^ and *d*^out(*v*)^ are the in-degree and out-degree of *v*, respectively. We return the number of edges that capture *τ*% of the total *f* (*u, v*) in the entire graph. RWR requires a damping factor α ∈ (0, 1] which is one minus the teleportation probability and a threshold *τ* ∈ (0, 1]. Default parameters are α = 0.85 and *τ* = 0.3. We take the negative log (*f* (*u, v*)) such that the predictions are ranked in increasing order.

#### All Pairs Shortest Paths (SP)

Finally, we also computed the shortest path from every source to every target and took the union of such paths as the reconstruction. If there are many tied paths between a source and a target, one path is arbitrarily chosen. The unranked set of edges in the shortest paths are considered the predicted pathway reconstruction.

### 2.3 Data

#### Interactome

We use PLNet_2_, a weighted, directed interactome constructed from both molecular interaction data and signaling pathway databases (including the database used for ground truth pathways) [16]. PLNet_2_ is weighted using an evidence-based Bayesian method introduced by RN [9] that assigns a high confidence to edges supported by experimental methods that successfully predict signaling interactions [16]. PLNet_2_ contains 17, 168 nodes (UniProtKB identifiers [27]) and 612, 516 directed edges, including 286, 520 physical interactions that are converted to bidirected edges.

#### Signaling Pathways

We consider a set of 29 signaling pathways from the NetPath database, a repository of cancer and immune related pathways [1]. Sources and targets are automatically detected for each of the pathways from curated lists of human receptors and transcriptional regulators (see [19] for more details). The Advanced Glycation End Products (AGE/RAGE) pathway, Inhibitor of Differentiation (ID) pathway, and Interleukin 11 (IL-11) pathway did not have an automatically-detected receptor and/or a transcriptional regulator, and were removed from consideration. Twenty-nine remaining pathways are listed in Table 2.

**Table 2.** NetPath pathways used in this study. Edges are taken as undirected for the assessment.

| Pathway<br>($n = 29$) | # Nodes | | | # Edges | |
| --- | --- | --- | --- | --- | --- |
|  | Total | \|S\| | \|T\| | Dir. | Undir. |
| $\alpha 6 \beta 4$Integrin | 66 | 7 | 3 | 223 | 116 |
| Androgen Receptor | 165 | 2 | 75 | 491 | 251 |
| BCR | 137 | 1 | 18 | 456 | 261 |
| BDNF | 72 | 5 | 4 | 139 | 76 |
| CRH | 24 | 2 | 8 | 54 | 27 |
| EGFR1 | 231 | 6 | 33 | 1456 | 756 |
| FSH | 19 | 2 | 1 | 30 | 18 |
| Hedgehog | 36 | 6 | 13 | 124 | 64 |
| IL1 | 43 | 3 | 5 | 178 | 93 |
| IL2 | 67 | 3 | 12 | 242 | 139 |
| IL3 | 70 | 2 | 9 | 176 | 97 |
| IL4 | 57 | 5 | 12 | 173 | 91 |
| IL5 | 30 | 2 | 3 | 71 | 36 |
| IL6 | 53 | 4 | 14 | 162 | 83 |
| IL7 | 18 | 2 | 3 | 52 | 28 |
| IL9 | 13 | 2 | 3 | 29 | 15 |
| Kit | 76 | 6 | 8 | 207 | 109 |
| Leptin | 55 | 3 | 15 | 135 | 74 |
| Notch | 74 | 4 | 27 | 255 | 154 |
| OSM | 37 | 5 | 12 | 82 | 42 |
| Prolactin | 68 | 4 | 10 | 199 | 103 |
| RANKL | 57 | 2 | 12 | 142 | 76 |
| TCR | 154 | 7 | 20 | 504 | 271 |
| TWEAK | 17 | 1 | 4 | 30 | 15 |
| TSLP | 7 | 1 | 2 | 14 | 7 |
| TSH | 48 | 2 | 6 | 90 | 47 |
| TGFβ | 209 | 5 | 77 | 863 | 452 |
| TNFα | 239 | 4 | 44 | 913 | 473 |
| Wnt | 106 | 14 | 14 | 428 | 220 |

### 2.4 Assessment

#### Precision and Recall

We evaluate reconstruction methods based on their precision as well as their recall. Let *G* = (*V, E*) be a potentially directed interactome. Let *P* = (*V*_*p*_, *E*_*p*_) be a pathway such that *V*_*p*_ ⊆ *V* and *E*_*p*_ ⊆ *E*. Let *H* = (*V*_*H*_, *E*_*H*_) be the output of some prediction method ℳ. Let *N* = (*V*_*N*_, *E*_*N*_) be a subgraph of *G* such that *N* is disjoint from *H* and such that the interactions and proteins in *N* are unlikely to be related to signaling. *N* is described in more detail in section 2.4.1. If *M* produces unranked predictions (as is the case for RN, BTB, PCSF, and SP), then the edge (or interaction) precision *p*_*E*_ and recall *r*_*E*_ of *M* for pathway *H* are

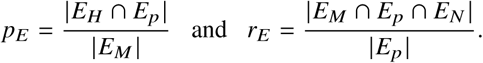

In the case that *E*_*N*_ ∪ *E*_*p*_ = *E* the equation for recall can be simplified to

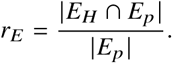

For ranked predictions we index precision and recall to a rank *i* such that *p*_*E*_(*i*) is the precision for the top *i* predictions. Thus the precision and recall for ranked methods (such as PL and RWR) comprise many points. Node (protein) precision and recall equations are the same *mutatis mutandis*.

We assessed the performance of reconstruction methods by way of their maximum *F*1 score, which we denote *F*_max_. Given a precision *p* and a recall *r*,

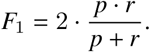

The *F*_max_ for a method is just the maximum *F*_1_ for that method. For unranked methods this is just the *F*_1_ score of their single precision-recall point.

#### 2.4.1 Negative Set

Because our task is not only to reconstruct *known* pathway interactions, but also to discover *unknown* interactions, we construct negative sets using sub-sampling. Following [19] we generate negative sets for each pathway by randomly selecting interactions which are not in the ground truth pathway at a rate of fifty negatives for every known interaction. However, the choice of sub-sampled negatives can affect a method’s performance in terms of precision. We designed a procedure to select a set of negatives for each pathway that is used for all precision-recall calculations for all reconstruction methods. For each pathway *P*, we compute the *F*_max_ for each of the six original methods (ℳ) using 50 distinct randomly sampled negative sets and place them in a 6*x*50 matrix *M*. Each column of *M* has the *F*_max_ scores for a given method across all 50 negative sets. We then create a new 6*x*50 matrix *N* such that

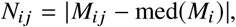

where med(**x**) is the median of a vector. Intuitively, the rows of *N* represent the differences from the median *F*_max_ for a given negative set for all methods. We sum the rows of *N* to get the aggregated difference from the median *F*_max_ for each negative set, and select the negative set that has the minimum of these values (reflecting the negative set that is the closest to the median across all methods). We do this for each pathway *P* in order to get reasonable negative sets across all pathways that do not advantage a particular method ℳ.

#### 2.4.2 Composites

In order to evaluate the overall performance of each prediction method across all pathways, we construct composite predictions and negative sets. Let *N*_*P*_ denote the set of negative edges for pathway *P* as described in section 2.4.1. We can create a composite negative set *N*_comp_ by taking a union over these edge sets, keeping track of the pathway for each negative edge:

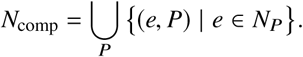

Likewise for a method ℳ with predictions *H* = (*V*_*P*, ℳ_, *E*_*P*, ℳ_) for a pathway *P*, let *H*_comp_ be the union over these edges:

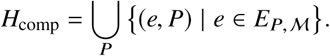

In the case of ranked predictions the edges of *H*_comp_ are then sorted such that a partial ordering is restored. The set of positives *P*_comp_ are constructed in a similar way from the ground truth pathways *P* = (*V*_*P*_, *E*_*P*_):

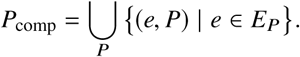

Precision, recall, and *F*_max_ can be calculated as described in Section 2.4, providing a general overall view of the performance of each method, though it privileges the performance of methods on larger pathways and obscures the heterogeneity of the performances on individual pathways.

## 3 Results

### 3.1 Reconstruction Methods Successfully Recover Proteins in Pathways

The Pathway Reconstruction Problem is solved by recovering both proteins and interactions that connect receptors to transcriptional regulators, and previous work has shown that recovering interactions is a challenging task for methods [19]. However, current pathway reconstruction methods are relatively successful at recovering pathway involved proteins. We evaluated the six original pathway reconstruction methods ℳ on their ability to recover the nodes in the Wnt pathway, subsampling negative nodes at a rate of 50 negatives to every positive (Figure 3). As has been previously shown [19], the methods maintain relatively high precision at larger values of recall — especially PL and RWR which offer ranked lists of candidate nodes (rather than a node set). Strikingly, if we ignore negative nodes that share an edge with a positive node, all methods jump to nearly perfect precision (starred methods in Figure 3). Thus, almost all predicted nodes for all methods are either in Wnt or one edge away from Wnt in *G*. These results indicate that using predicted nodes from existing pathway reconstruction methods may be able to leverage reliable information to predict protein interactions.

**Fig 1.**
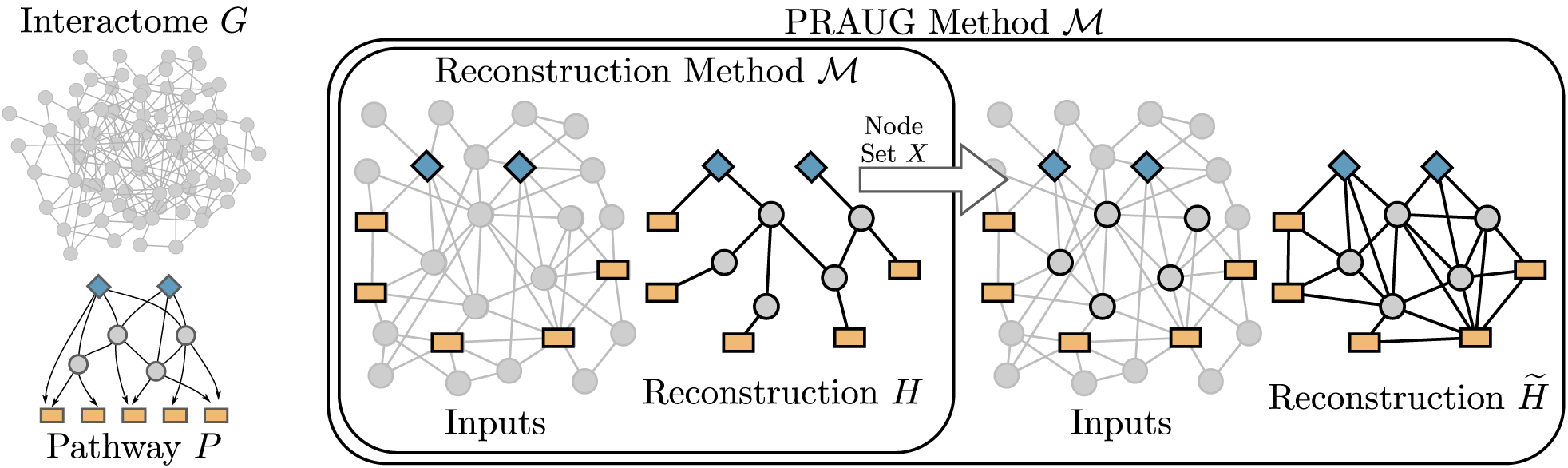
Overview of PRAUG. Pathway reconstruction methods take as input an interactome *G* and a pathway comprised of sources (blue diamonds) and targets (orange rectangles). Given a pathway reconstruction method ℳ, PRAUG defines a new method 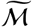 that calls ℳ and performs a traversal on the resultant node set.

**Fig 2.**
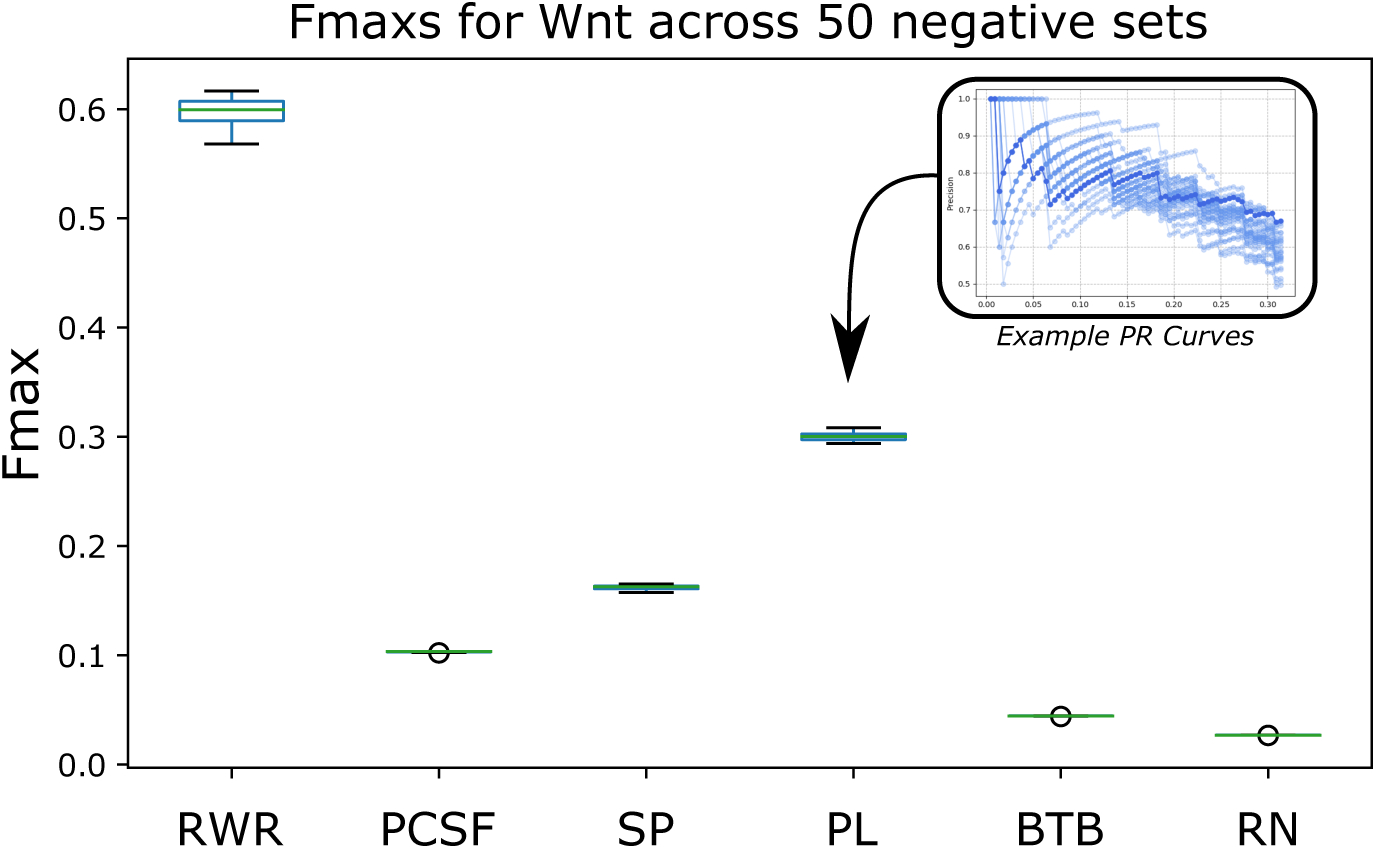
*F*_max_ scores for each method across 50 distinct sub-sampled negative sets for the Wnt pathway. Inset shows example precision-recall curves for PathLinker that are used to compute the *F*_max_ distribution.

**Fig 3.**
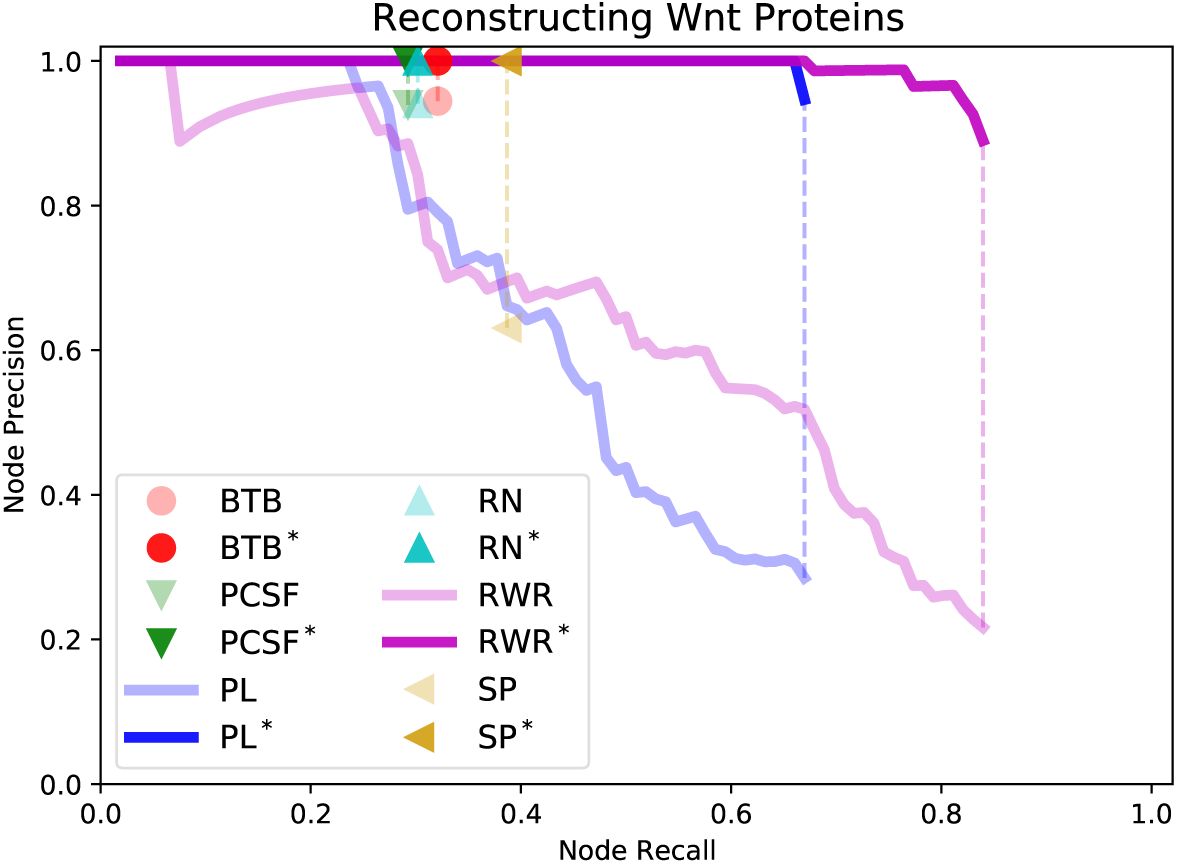
Existing methods recover proteins with high precision. Asterisks denote that negative nodes connected to a positive are ignored. (PL was run with *k* = 5000 and RWR was run with *τ* = 0.5 for illustration purposes.)

#### 3.1.1 Prediction Overlaps

In order to assess the similarity of different reconstruction methods we computed the Jaccard overlap of the proteins they predicted across all pathways using default parameters. Given the interactome *G* = (*V, E*), let *A* ⊆*V, B* ⊆ *V*, be two sets of nodes. Then the asymmetric Jaccard overlap of *A* with *B* is the percentage of elements of *A* which are also in *B*. We computed the Jaccard overlap of composite predictions *H*_comp_ for the six methods across all 29 pathways (Figure 4). The algorithms, which are quite heterogeneous, have a considerable amount of agreement in their reconstructed proteins, ranging from Jaccard indices of 0.18 (PCSF with RWR) to 0.95 (BTB with SP).

**Fig 4.**
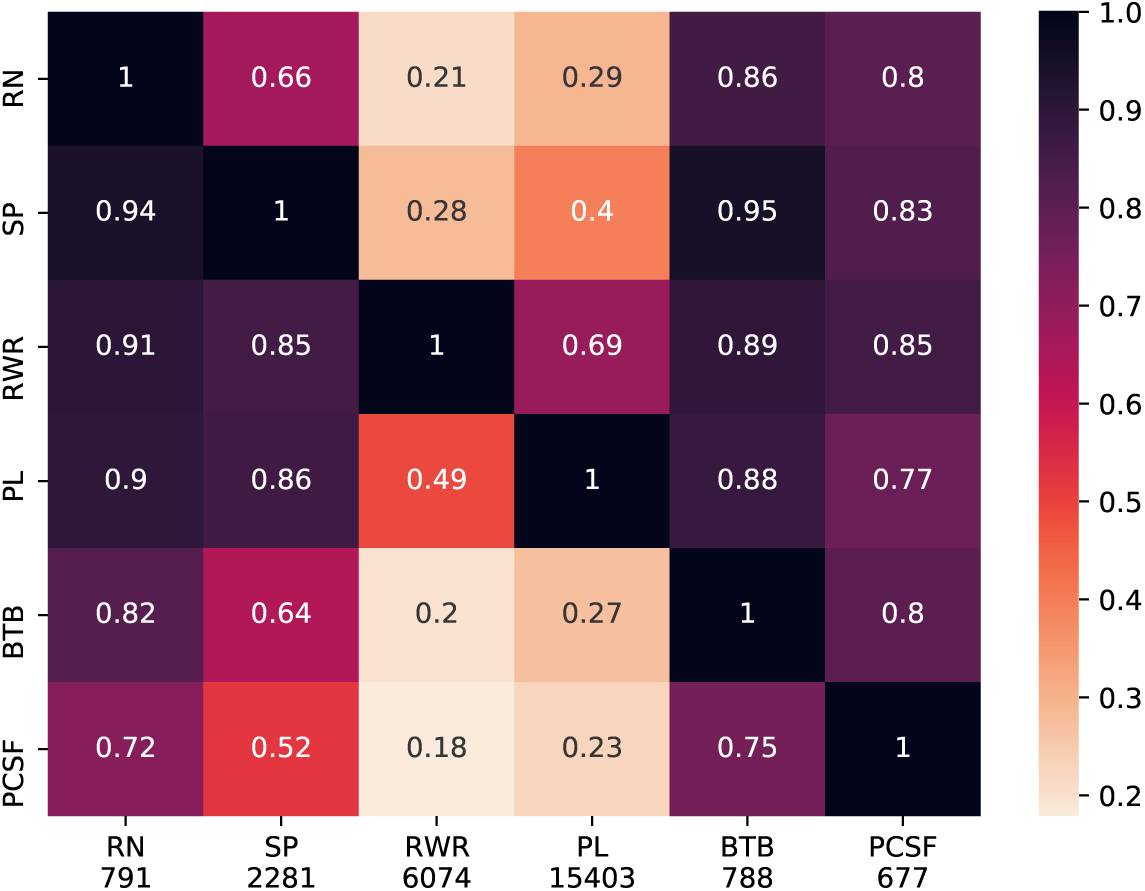
Jaccard overlap of nodes predicted by pathway reconstruction methods, normalized by the columns. Number of predicted interactions given below x-axis labels.

### 3.2 PRAUG Improves Upon Reconstruction Methods

We have shown that reconstruction methods can recover the proteins in a pathway; however recovering the interactions is more challenging. We compared each reconstruction method to PRAUG in terms of precision and recall across all 29 signaling pathways (Figure 5). Each panel compares the performance of a pathway reconstruction method ℳ from Section 2.2 to 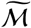, using default values and taking care to keep the negative set consistent for each pathway (Section 2.4.1). While we note that there are differences among reconstruction methods, we did not optimize the parameters for each method with respect to the Pathway Reconstruction Problem, so we do not compare un-augmented methods against one another.

**Fig 5.**
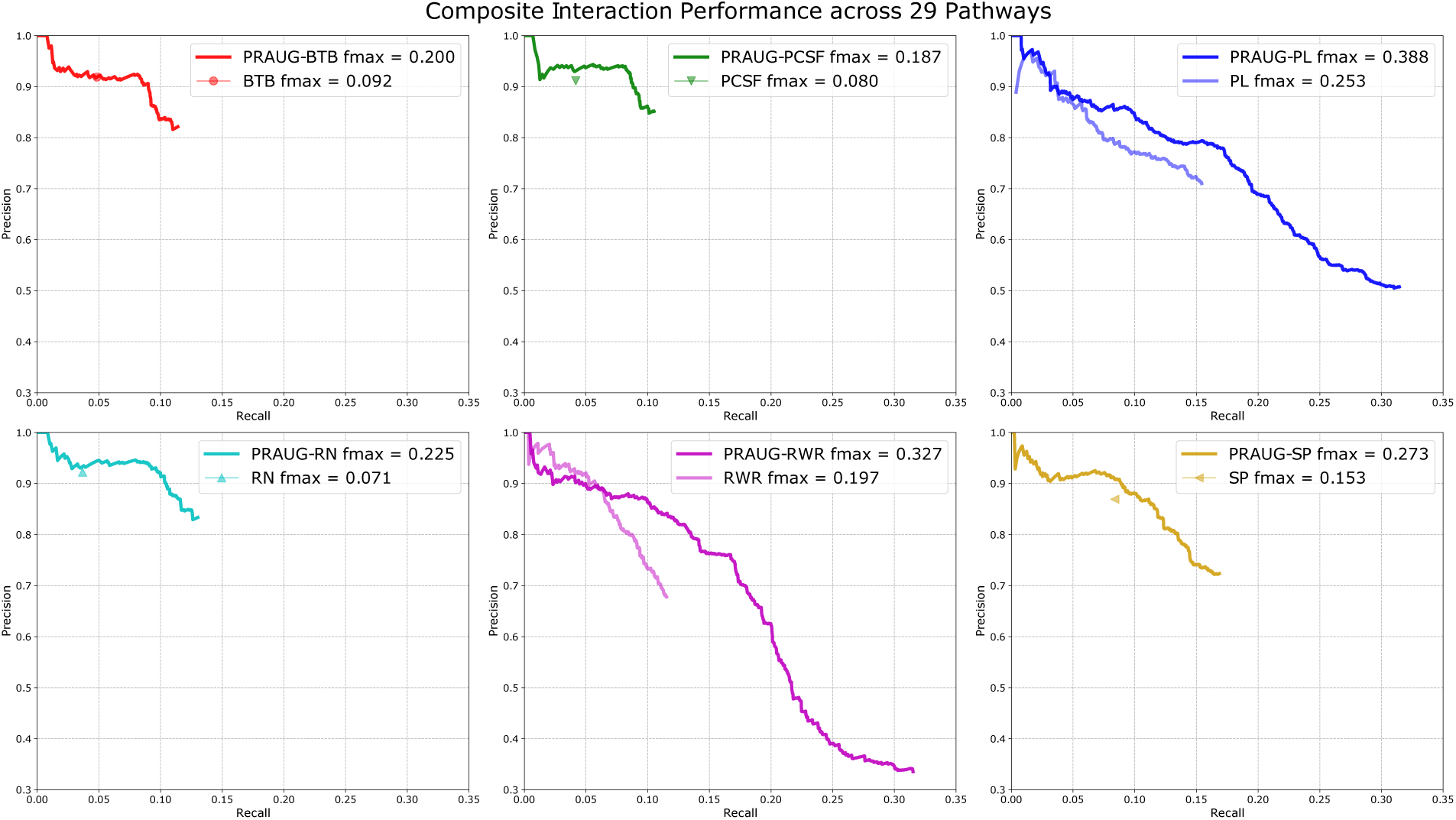
Performance of method ℳ and PRAUG-transformed 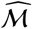across all 29 pathways.

In every case, the PRAUG method 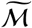’s *F*_max_ value is larger than the original method ℳ’s *F*_max_ in terms of recovering the interactions, outperforming ℳ by an average of 89% (Figure 5). PRAUG gains this large improvement in *F*_max_ by improving recall while sacrificing sparingly with respect to precision. While precision at maximum recall is lower across the board for augmented methods as compared to their counterparts, precision for 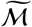 is equal to or greater than the precision of ℳ at the top recall of ℳ.

The methods ℳ that produce unranked predictions (BTB, PCSF, RN, and SP) are single points in Figure 5, and the PRAUG-augmented counterparts of these methods maintain the precision of ℳ for larger values of recall. However, many exhibit a plateau of precision after which precision drops off rapidly with increased recall (for example, precision remains nearly constant for PRAUG-BTB from recall values of 0.0125 to 0.08). The PRAUG counterparts of the ranked methods (PL and RWR) do not exhibit this behavior. Despite the differences between the unranked and ranked methods, the key result is that the PRAUG counterparts 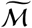 of each pathway reconstruction method outperforms ℳ in *F*_max_, recall, and often precision.

### 3.3 Robustness Analysis and Benchmarking

PRAUG is a straightforward traversal-based algorithm, and it may not be immediately clear why this approach can be so successful. In this section we justify PRAUG’s depth-first traversal, provide empirical upper bounds on the Pathway Reconstruction Problem given methods which assume a traversal paradigm, and illustrate the effect of parameter selection on PRAUG methods compared to their original counterparts.

#### 3.3.1 Choice of Traversal

Step 4 of a PRAUG-augmented pathway reconstruction method is to perform a depth first traversal from the sources of a pathway ignoring edge weights (Section 2.1). We considered three alternatives to PRAUG’s unweighted depth-first traversal:

**PRAUG-BFS:** Perform an unweighted breadth-first traversal on *G* starting from *σ*, taking only nodes in *X*.

**PRAUG-WEIGHTED:** Perform a weighted depth-first traversal on *G* starting from *σ*, taking only nodes in *X*.

**PRAUG-BFS-WEIGHTED:** Perform a weighted breadth-first traversal on *G* starting from *σ*, taking only nodes in *X*.

In all cases, the method terminates when no new edge can be traversed from *σ* to nodes in *X*. Thus, all of these methods recover the exact same interactions but traverse them in a different order. The choice of traversal or edge weighting has little effect on the precision and recall, as is illustrated for PL and RWR on the Wnt pathway (Figure 6). When there is a difference, using breadth first traversal performs worse than using depth first traversal, and weighting has a negligible effect on the performance of the augmented methods.

**Fig 6.**
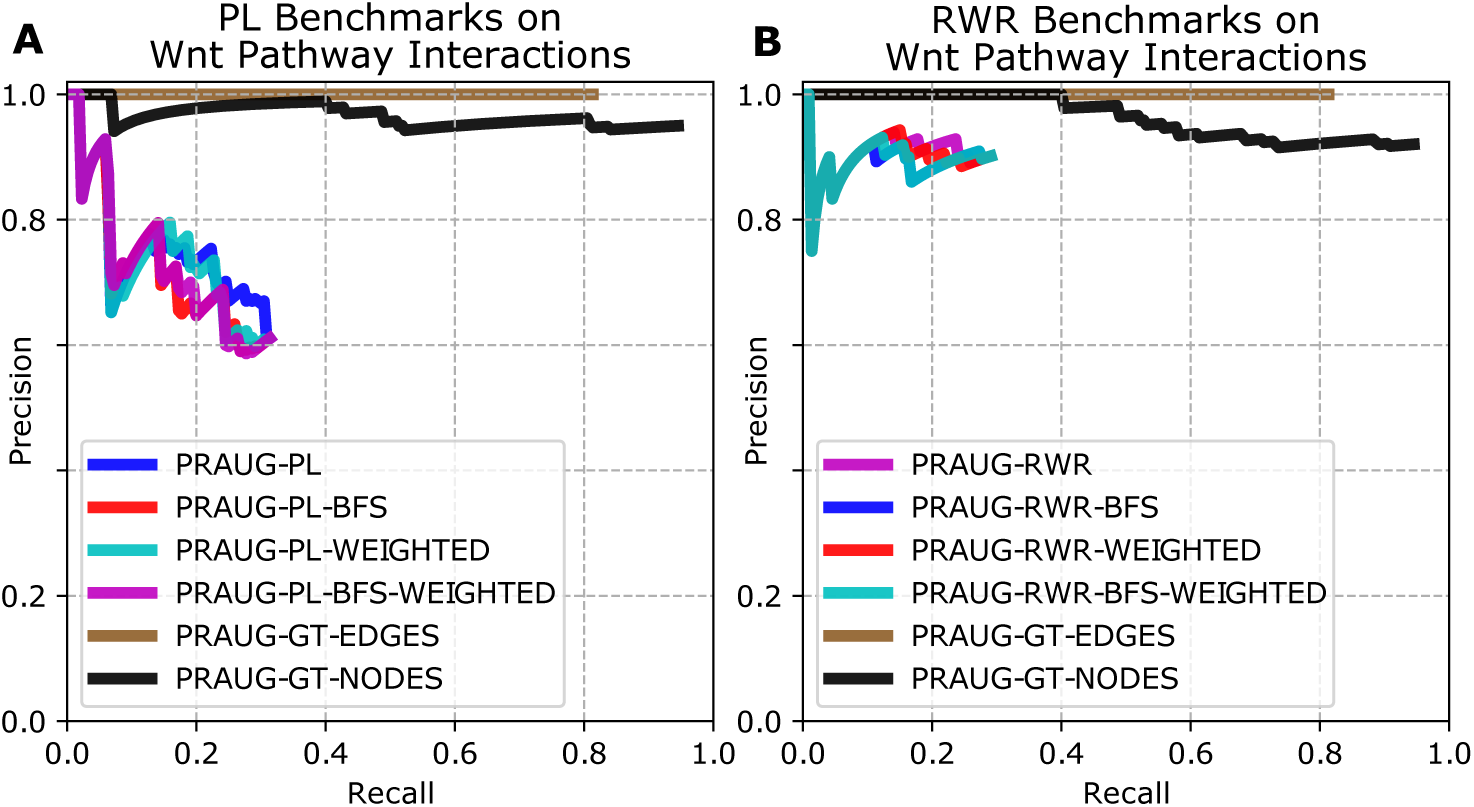
PRAUG compared to breadth-first search, using a weighted interactome, and using ground truth (GT) nodes and edges as inputs for (A) PL and (B) RWR.

#### 3.3.2 Ground Truth as Upper Bounds

We also investigated how well PRAUG could recover the pathways when given the ground truth nodes and interactions, which serve as an upper bound on a traversal-like performance. We first define an oracle pathway reconstruction method GT which, given an interactome *G*, a source set *S*, and a target set *T*, produces the unique ground truth signaling pathway *P* = (*V*_*p*_, *E*_*p*_) such that *S* ⊆ *V*_*p*_ and *T* ⊆ *V*_*p*_.^1^ Of course, just because we can define something doesn’t mean we can build it. If we knew how to realize GT — an algorithm which produces the ground truth pathway by definition — then we would doubtlessly be publishing a paper on that instead. In lieu of such good fortune we provide our implementation of GT with the actual pathway *P* to be produced. Thus, our implementation of GT does not meaningfully solve the pathway reconstruction problem. However, GT provides the complete set of ground truth nodes to its PRAUG-augmented counterpart:

**PRAUG-GT-NODES:** Perform an unweighted depth-first traversal on *G* starting from *σ*, taking only nodes in *V*_*p*_.

PRAUG-GT-NODES informs us how well a PRAUG-augmented method could perform if a pathway reconstruction method ℳ perfectly reconstructed the proteins involved in a given pathway. In this first upper-bound, PRAUG-GT-NODES provides the precision and recall of interactions assuming (a) perfect node precision and recall and (b) we begin the traversal from the sources.

We can ask a similar question about PRAUG’s performance when we have the ground truth interactions available for traversal. This requires augmenting the definition of Step 4 of PRAUG as follows:

**PRAUG-GT-EDGES:** Perform an unweighted depth-first traversal on *G* starting from *σ*, taking only *edges* from *E*_*p*_.

PRAUG-GT-EDGES differs from PRAUG because it traverses edges rather than nodes. PRAUG-GT-EDGES is guaranteed to maintain perfect precision and marks the upper bound on recall for a traversal based method starting from the sources which only makes predictions from the annotated ground truth.

In Figure 6, PRAUG-GT-NODES are shown in black and PRAUG-GT-EDGES are shown in brown. The ground truth Wnt pathway is the same in both panels A and B, and the only difference between panels is the subsampled negative set involved in the calculation. PRAUG-GT-NODES achieves nearly total recall while maintaining nearly perfect precision for Wnt, which is representative of PRAUG-GT-NODES generally. As expected, PRAUG-GT-EDGES maintains perfect precision up to much larger values of recall than any other pathway reconstruction method. Interestingly, PRAUG-GT-NODES achieves higher recall than PRAUG-GT-EDGES, because there are some incorrect interactions that allow the traversal to reach more positive interactions when only considering nodes.

#### 3.3.3 Parameter Sweeps

Several of the reconstruction methods considered in this paper take additional inputs beyond an interactome, source nodes, and target nodes (Table 1). These additional parameters substantially affect the behavior of the methods. In order to assess how additional parameters affected the performance of both original methods ℳ as well as their corresponding augmented methods 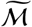, we performed parameter sweeps for both PRAUG-PL and PRAUG-RWR (Figure 7). PL and RWR both have parameters which determine the number of edges in the reconstructed pathway. In the case of PL, *k* denotes the number of paths returned.^2^ In the case of RWR, *τ* marks the percentage of total edge flux in the interactome to return. Thus for PL (resp. RWR) lower *k* (resp. *τ*) values produce truncated precision recall curves of higher *k* (resp. *τ*) values.

**Fig 7.**
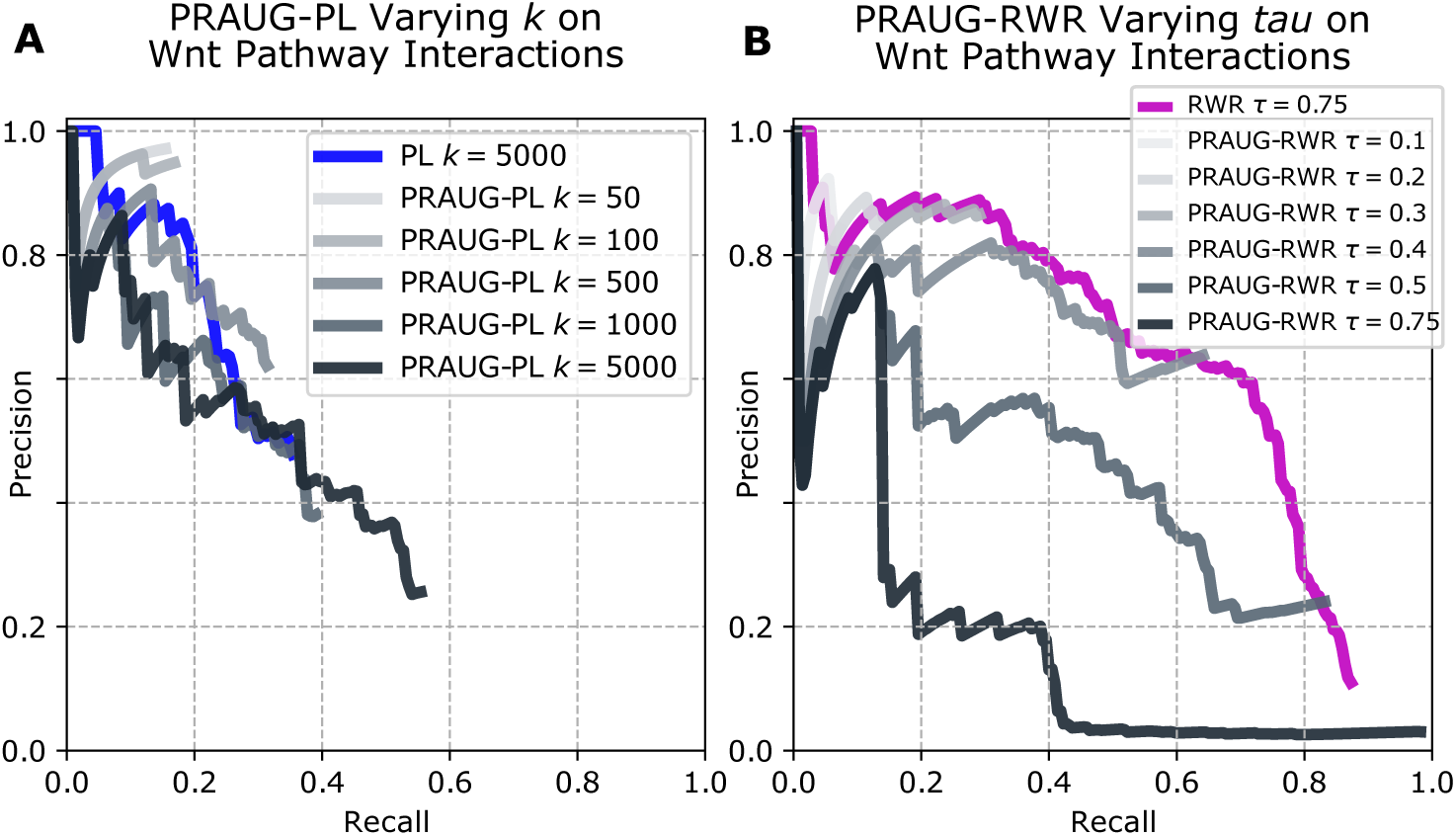
PRAUG performance when (A) varying *k* for PL and (B) varying *τ* for RWR. PL (blue) and RWR (magenta) are shown for the largest parameter, since all smaller parameters simply truncate this curve.

The same does not hold for the PRAUG-augmented counterparts to PL and RWR. Because the ranking of edges in a PRAUG-augmented method is given by their order in the traversal, the introduction of new nodes (by way of parameter change) to the traversal leads to markedly different precision recall curves (Figure 7). Further, PRAUG-augmented methods are highly sensitive to parameter changes. In general large parameters lead to degraded precision at low values of recall, but recover a large portion of positive interactions. This can be seen for instance in PRAUG-RWR *τ* = 0.75 in Figure 7B. Thus, optimal parameters for an unaugmented method may deviate significantly from the optimal parameters for it’s PRAUG-augmented counterpart.

### 3.4 Case Studies

To illustrate the benefits of pathway reconstructions augmented by PRAUG, we focus on two well-studied signaling pathways: Wnt and Notch. For each pathway, we show the single-pathway precision and recall curves for all methods ℳ and 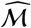, as well as PRAUG-GT-NODES and PRAUG-GT-EDGES. We also consider the first 500 edges predicted by each of the six PRAUG methods, and we visualize the interactions that are supported by all six methods. It is important to note that PRAUG ranks interactions according to the order in which edges are traversed, which does not necessarily imply confidence in the interaction. While the choice of using the first 500 interactions is arbitrary, they provide a representative set of interactions from the traversal and produce a network that connects most of the receptors to transcriptional regulators.

### 3.5 The Wnt Signaling Pathway

The canonical Wnt signaling pathway regulates a variety of developmental events during embryogenesis and helps maintain tissue homeostasis [28]. The non-canonical Wnt signaling pathway, which is not mediated through the canonical Frizzled receptors or *β*-catenin, also regulates cell movement and tissue polarity [1]. The combination of canonical and non-canonical Wnt signaling makes this a good case study for PRAUG reconstructions. Pathway reconstruction methods that return a single subgraph tend to have high precision but quite low recall (Figure 8A). PL and RWR extend this recall while maintaining relatively high precision. which has been previously shown [19]). The PRAUG methods are able to improve both precision and recall for all methods except for PL, in which it only improves recall (Figure 8A). PRAUG-GT-EDGES and PRAUG-GT-NODES are described in Section 3.3.2 and are provided here for benchmark purposes.

**Fig 8.**
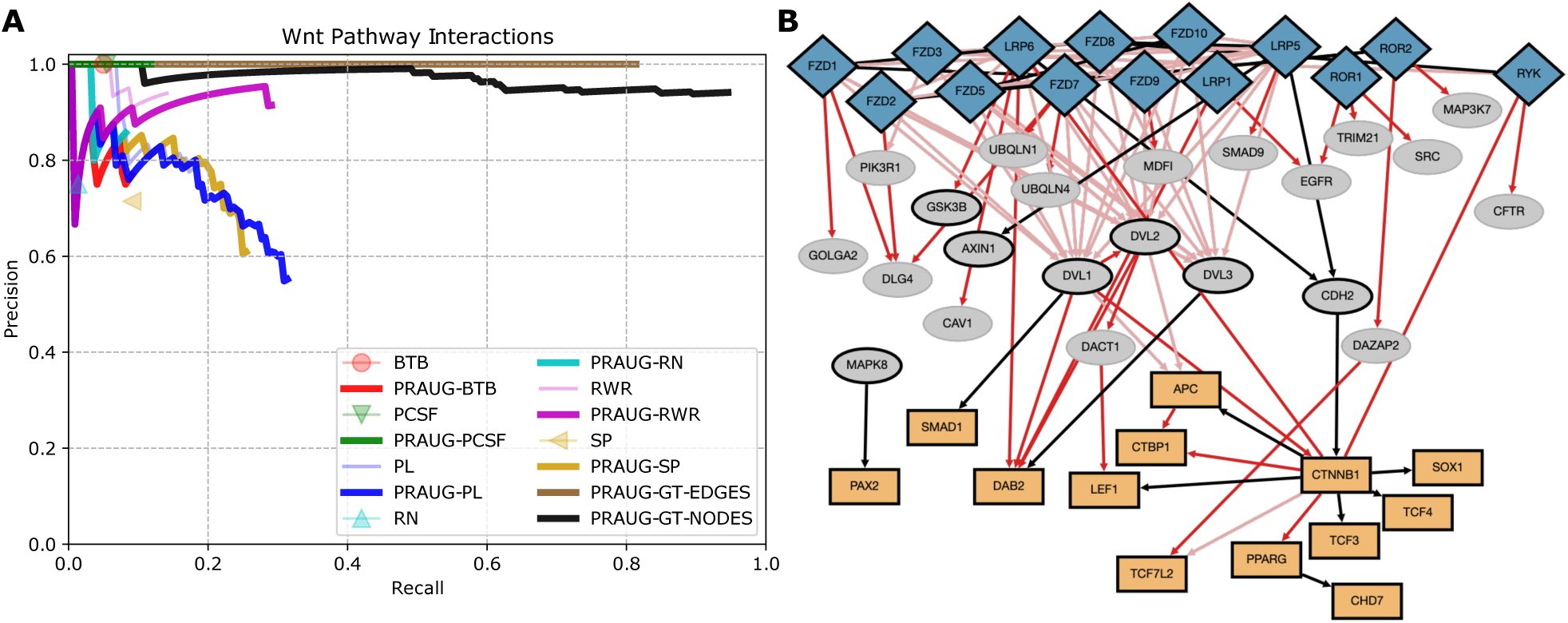
Precision recall curve for the Wnt signaling pathway (A), and the network comprised of the top interactions predicted by the PRAUG reconstructions (B). Blue diamonds are receptors (sources *S*) and orange rectangles are transcriptional regulators (targets *T*). Black nodes and edges appear in the Wnt NetPath pathway; pink edges appear in some KEGG pathway, and red edges do not appear in NetPath or KEGG databases.

The interactions that appear in the first 500 interactions of all PRAUG methods (Figure 8B) are largely consistent with the pathway reconstruction reported by the first 200 paths from PathLinker [19]. The Wnt network includes CFTR, a chloride ion transporter that interacts with non-canonical RYK receptor, which has been shown to be involved in Wnt signaling via a subset of the Wnt ligands [19]. Many of the 22 intermediate gray nodes have previously been implicated in Wnt pathway crosstalk, including the MAP Kinases, SRC, EGFR, PII3KR1, and SMAD9 [19, 29]. There are also known ubiquitination proteins (UBQLN1, UBQLN4, and TRIM21) which are general proteins and may not be specific to Wnt signaling [27]. Other non-positives have documented roles in Wnt signaling despite not being present in NetPath: for example, DACT1 interacts through Disheveled (DVL) family proteins and MDFI regulates TCF family transcription factors [27]. Further, proteins such as DLG4 and DAZAP2 are not directly associated with Wnt signaling, and may offer new hypotheses for follow-up investigation. DLG4 is particularly promising, as it is involved in cell-cell adhesion (one of Wnt’s classic roles) but is known to interact with the NMDA receptor in the brain [27].

In addition to the nodes within the Wnt network, the interactions themselves also help put the reconstructions in context. Only a handful of the interactions in the Wnt network appear in the NetPath Wnt pathway (black edges in Figure 8B). The remaining edges are either negatives or ignored due to subsampling in the precision-recall plots. However, 66% of these edges appear as interactions in some KEGG signaling pathway (shown in pink), indicating that they are signaling interactions. Further, some of the edges that do not appear in NetPath or KEGG (red edges) connect proteins in NetPath’s Wnt signaling (e.g. DVL1-DVL2 and LRP6-GSK3B), which may be known interactions that have been missed by signaling pathways. In this way, pathway reconstruction methods that focus on recovering interactions have the potential for generating more mechanistic-driven hypotheses.

### 3.6 The Notch Signaling Pathway

The Notch signaling pathway regulates fundamental processes such as morphogenesis, cell differentiation and cell-fate determination, proliferation, and cell death, and is highly conserved across metazoans [1, 30]. Notch is a unique case study for the Pathway Reconstruction Problem because its primary receptors function both as cell surface receptors and nuclear transcription factors, seemingly contradicting our assumptions that a signaling pathway’s receptors and TRs are mutually exclusive. However, most pathway reconstruction methods recover Notch signaling with high precision at low recall, and PRAUG methods typically improve recall while remaining close the original method’s precision (Figure 9A). Unlike Wnt, PRAUG-GT-EDGES for Notch only recovers about half of the positive interactions, but PRAUG-GT-NODES boasts a recall greater than 0.8.

**Fig 9.**
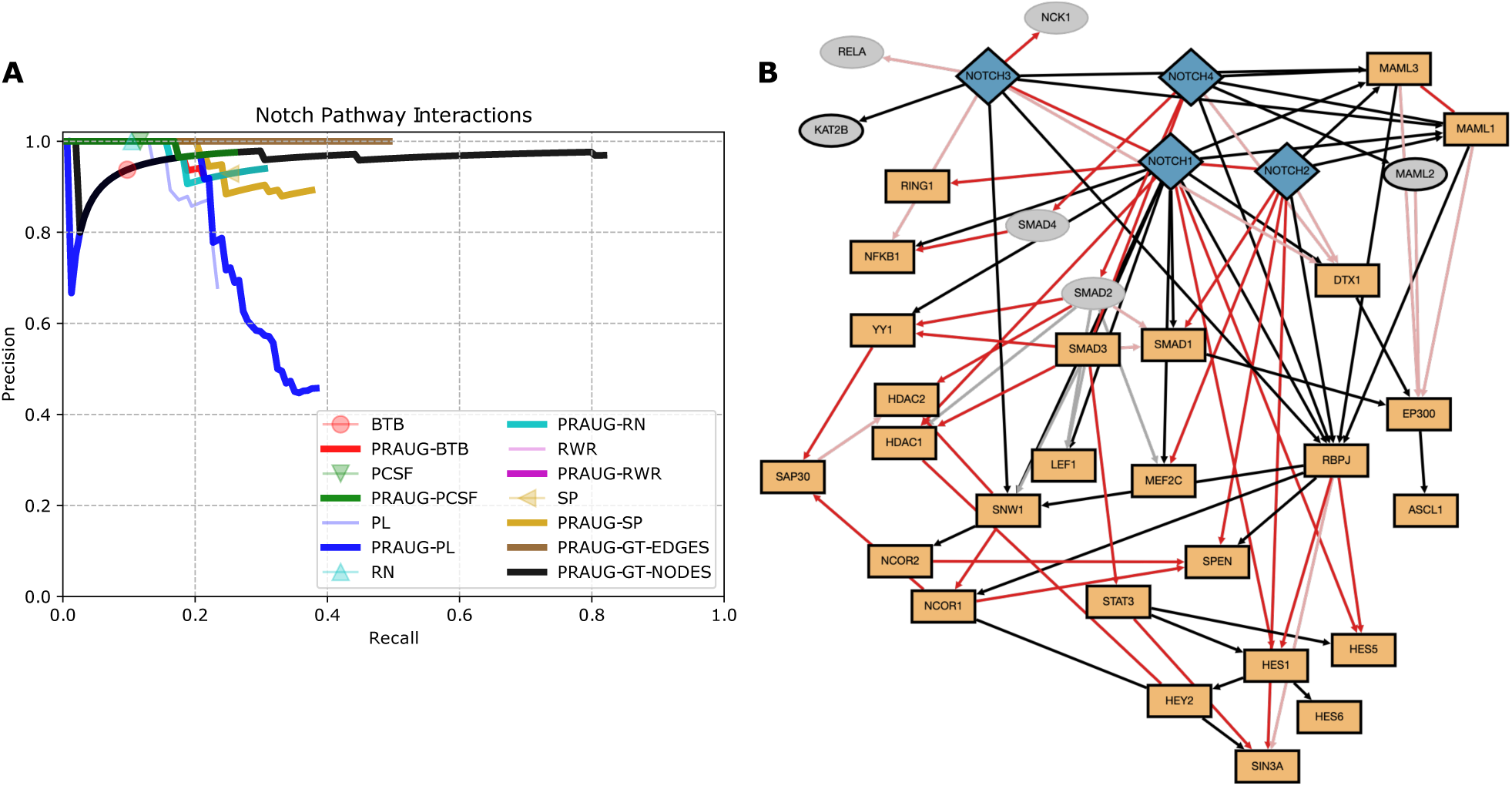
Precision recall curve for the Notch signaling pathway (A), and the network comprised of the top interactions predicted by the PRAUG reconstructions (B). Edges and nodes are colored as described in Figure 8; and gray edges appear in some NetPath pathway other than Wnt.

When we examine the interactions recovered in the first 500 interactions of all six PRAUG methods, we see that the majority of the interactions connect receptors to TRs, highlighting the nuclear role of Notch family proteins (Figure 9B). One of the six intermediate proteins is a member of the MAML transcriptional coactivator, and two are SMAD proteins, which along with NFκB are known to associate with the Notch intracellular domain [30]. Five of the 53 non-positive interactions appear in NetPath’s TGF-*β* Receptor pathway [1], and another twelve interactions appear in KEGG as members of Notch signaling (e.g. edges from MAML proteins to EP300) or Th1 and Th2 cell differentiation (e.g. edges from Notch3 to RELA and NFκB1, which co-complex to form NFκB), of which Notch is known to initiate [31]. While two thirds of non-positive interactions do not appear in NetPath or KEGG (shown in red), they connect proteins that are in the NetPath Notch pathway, suggesting that these may be missed interactions in NetPath. Further, the connection from Notch3 to NCK1, an adaptor protein which associates with certain growth factor receptors, has recently been shown to maintain Notch signaling by preventing processes induced by Notch inhibition, and provides a promising hypothesis to test [32].

## 4 Discussion

We present PRAUG, a higher-order algorithm for augmenting pathway reconstruction methods which consistently recovers interactions from ground truth pathways better than the original methods. We have shown that this framework is highly generalizable, improving reconstructions for six pathway reconstruction algorithms that include shortest-paths (PL, BTB, SP), network flow (RN), random walk (RWR), and Steiner tree (PCSF) approaches. We demonstrate an improved *F*_max_ when PRAUG is applied to all of these methods across 29 signaling pathways (Figure 5), which cover diverse pathways with respect to both size (e.g. number of nodes and interactions in Table 2) as well as mechanism (immune signaling pathways, cancer hallmark pathways, and hormone signaling pathways). We illustrate PRAUG’s power in generating plausible hypotheses for follow-up validation by analyzing interactions predicted by all PRAUG method predictions for Wnt and Notch (Figures 8 and 9).

PRAUG does not present a new pathway reconstruction algorithm, but rather a paradigm for improving other algorithms designed to reconstruct pathways. The foundational observation of our method is that reconstructing the nodes in a pathway is a relatively easy task at which many existing algorithms excel (Figure 3). In particular, The performance of PRAUG-GT-NODES demonstrates that PRAUG can produce substantially better pathway reconstruction methods if it is provided with a method which reliably produces correct node sets. Even when limiting the inputs to ground truth edges, PRAUG-GT-EDGES achieves recall much higher than current algorithms. Every algorithm that we used for this paper attempts to explicitly reconstruct the *interactions* of a pathway; further work to improve protein reconstructions may lead us closer to these empirical upper bounds.

There are variants of the Pathway Reconstruction Problem, such as a “curation-aware” pathway reconstruction that uses all proteins and interactions from pathway databases as input [33], and pathway reconstructions that involve changing the underlying interactome [34]. Considering a traversal-based approach for these formulations may prove similarly useful. Further, PRAUG offers a multitude of opportunities for defining a “pathway reconstruction method” ℳ – for example, we could focus on gene ontology node sets or experimentally-derived genes as the node set *X* for PRAUG traversal. In this way, PRAUG may be able to recover interactions from methods which explicitly produce node reconstructions.

Of the two pathway reconstruction methods with ranked interactions, we were surprised by the dramatic performance of RWR. For large values of *τ*, RWR achieves precision of 0.8 at a recall of 0.4, blowing away all other non-PRAUG methods (magenta curve in Figure 7B). This method, which was inspired by TieDIE [11], performs a random walk from sources, then another random walk on a reversed graph from targets, and combines the edge flux scores for every edge. This random walk was different than the one presented in the PathLinker paper [19], which performed a random walk with restarts only from the sources. Here, RWR shows significant promise for improving pathway reconstructions, and is an improvement over prior algorithms in its own right.

Finally, we note that PRAUG is quite straightforward. Despite its simplicity, we described some subtleties to this method. For example, an optimal choice of parameters for a method ℳ may not be optimal for the PRAUG version 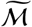 (Figure 7). In general, we found that the larger the reconstruction returned by ℳ, the worse the PRAUG augmented method performed. In essence, if the pathway reconstruction returned a large subset of the nodes in *G*, then PRAUG would return an induced subgraph of the nodes, which is bound to have many negative interactions. We do not explicitly compare pathway reconstruction methods to each other, as more careful selection of parameters for each method ℳ and PRAUG method 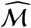 would be necessary.

PRAUG shows great potential for improving upon existing pathway reconstruction methods. We have shown six applications of PRAUG using extant reconstruction methods, and demonstrates promise for improving untested algorithms. PRAUG brings us one step closer to practical pathway predictions for emerging and under-studied signaling pathways.

### Data Availability

All code and datasets are available on https://github.com/TobiasRubel/Pathway-Reconstruction-Tools

Users can use GraphSpace [35] to explore the networks in Figures 8B and 9B:

### Wnt Network

http://graphspace.org/graphs/28928.

### Notch Network

http://graphspace.org/graphs/28927.

## Acknowledgments

This work was supported by the National Science Foundation (BIO-ABI #1750981 to AR).

## Footnotes

1 The oracle GT is an element of the set of all pathway reconstruction methods 𝕄.

2 Though note that new paths may use no new edges. Thus *k* neither fully nor uniquely determines the number of edges in the reconstructed pathway.

